# DIRseq: a method for predicting drug-interacting residues of intrinsically disordered proteins from sequences

**DOI:** 10.1101/2025.05.02.651562

**Authors:** Matthew MacAinsh, Sanbo Qin, Huan-Xiang Zhou

## Abstract

Intrinsically disordered proteins (IDPs) are now well-recognized as drug targets. Identifying drug-interacting residues is valuable for both optimizing compounds and elucidating the mechanism of action. Currently, NMR chemical shift perturbation and all-atom molecular dynamics (MD) simulations are the primary tools for this purpose. Here we present DIRseq, a fast method for predicting drug-interacting residues from the amino-acid sequence. All residues contribute to the propensity of a particular residue to be drug-interacting; the contributing factor of each residue has an amplitude that is determined by its amino-acid type and attenuates with increasing sequence distance from the particular residue. DIRseq predictions match well with drug-interacting residues identified by NMR chemical shift perturbation and other methods, including residues L_22_WK_24_ and Q_52_WFT_55_ in the tumor suppressor protein p53. These successes augur well for deciphering the sequence code for IDP-drug binding. DIRseq is available as a web server at https://zhougroup-uic.github.io/DIRseq/ and has many applications, such as virtual screening against IDPs and designing IDP fragments for in-depth NMR and MD studies.

## Introduction

Intrinsically disordered proteins (IDPs) are now recognized as important drug targets ^1-4^. For structured protein targets, a crucial step is identifying the drug-binding pocket. Although an IDP can be locked into a specific conformation by a drug molecule in rare cases ^5^, the prevailing scenario is that the protein remains disordered upon drug binding ^6-14^. Consequently, the IDP-drug complex typically samples a vast conformational space, and the drug molecule only exhibits preferences, rather than exclusiveness, for interacting with subsets of residues ^8,12,13^. Such drug-interacting residues, akin to binding pockets in structured proteins, are key to optimizing compounds ^13,15,16^ and elucidating the mechanism of action ^6,9,12,15,16^.

NMR chemical shift perturbation (CSP) is the best experimental method for identifying drug-interacting residues and has been applied to many IDPs ^7,9,11-26^. Aromatic residues are frequently among drug-interacting residues ^12,13,15,16,18,20,24^. This is understandable as drug molecules typically are rich in aromatic rings (Figure 1 and Table S1), which can form π-π interactions with aromatic residues. As recently pointed out by Heller et al. ^14^, CSPs of IDPs elicited by drug binding can be small enough to fall within the spectral resolution of NMR spectroscopy, therefore making it difficult to unequivocally identify drug-interacting residues. The small magnitude of CSPs may arise because the drug does not have a strong preference for interacting with any residues. Another scenario may be that drug binding induces a conformational shift such as secondary structure formation or even partial folding ^9,11,13^, so CSPs spread from the directly interacting residues to the rest of the IDP, adding to the difficulty in identifying drug-interacting residues.

**Figure 1.**
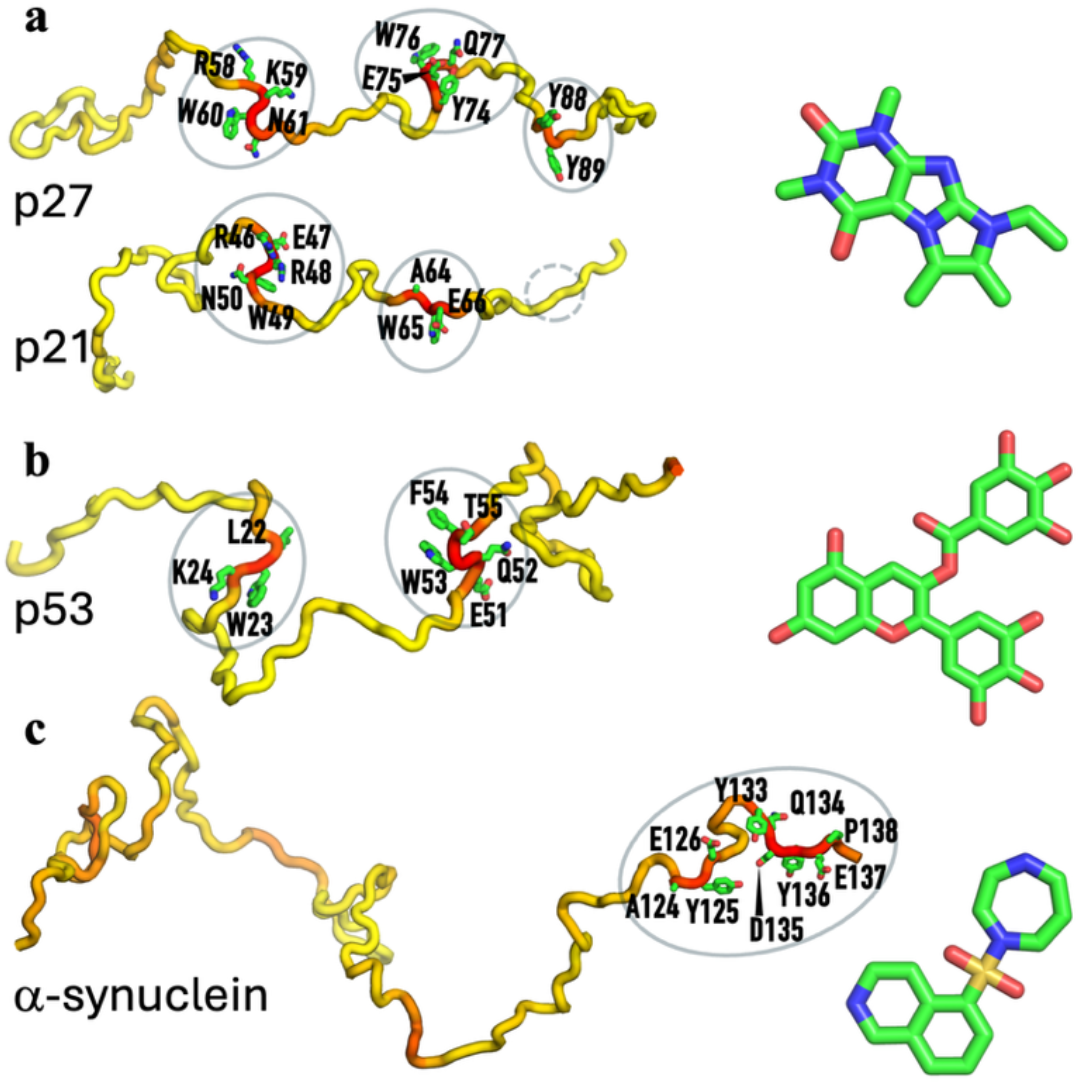
Four IDPs and drugs that bind to them. (a) p27, p21, and SJ403. (b) p53 and EGCG. (c) α-synuclein and Fasudil. For each IDP, the DIRseq propensities are rendered by a color spectrum from yellow for low values to red for high values. Predicted drug-interacting residues are shown with sidechains rendered in stick.

All-atom molecular dynamics (MD) simulations have presented atomic details in many IDP-drug systems ^8,10,12,13,16,24,27-31^. These simulation studies have highlighted the frequent engagement of drug molecules with aromatic residues, particularly in π-π interactions ^12,13,16,24,28-31^. MD simulations also revealed the emergence of a compact sub-population of the N-terminal disordered region of the tumor suppressor protein p53 upon binding the drug epigallocatechin-3-gallate (EGCG) ^12^ and reduced solvent exposure of hydrophobic residues in the 42-residue amyloid-β peptide (Aβ42) upon binding the drug 10074-G5 ^10^. However, MD simulations of IDPs still suffer from the perennial issues of inaccurate force fields and insufficient sampling, which are exacerbated when drug molecules are included. For example, two replicate simulations may identify different drug-interacting residues ^32^.

Here we present a sequence-based method, DIRseq, as a complement to NMR CSP and MD simulations. This method was motivated by our observation that drug-interacting residues seem to overlap with residues exhibiting elevated transverse relaxation rates (*R*_2_) ^33,34^, as exemplified by the C-terminal 20 residues of α-synuclein ^13^. Elevated *R*_2_ in IDPs is caused by either local inter-residue interactions or residual secondary structure ^35^. Based on this understanding, we developed a sequence-based method, SeqDYN, to predict *R*_2_ of IDPs ^33^. As suggested previously ^34^, the propensities of residues to form intramolecular interactions and therefore elevate *R*_2_ should be similar to those for forming intermolecular interactions with drug molecules. DIRseq is therefore an adaptation of SeqDYN. DIRseq predictions match well with drug-interacting residues identified by NMR chemical shift perturbation and other methods, including residues L_22_WK_24_ and Q_52_WFT_55_ in p53 and C-terminal residues in α-synuclein. DIRseq is available as a web server at https://zhougroup-uic.github.io/DIRseq/ and has many applications.

## Results

### Retooling of SeqDYN into DIRseq

SeqDYN ^33^ predicts the *R*_2_ value of residue *n* by accumulating a contributing factor *f*(*i*; *n*) from each residue *i*:

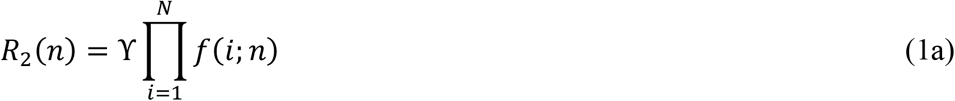

where *N* is the total number of residues in the IDP and ϒ is a uniform scale parameter. The contributing factor *f*(*i*; *n*) has an amplitude *q*(*i*) that is determined by the amino-acid type of residue *i* and attenuates with increasing sequence distance, *s* = |*i* − *n*|, from residue *n*:

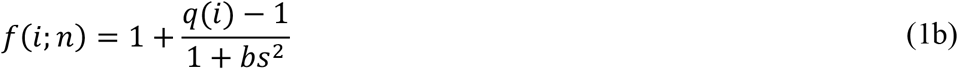

The 20 *q* parameters (one for each amino-acid type) and *b* were trained on the measured *R*_2_ values for 45 IDPs. The original *q* values (Figure S1a) show that aromatic (Trp, Tyr, His, and Phe), Arg, and long aliphatic (Ile and Leu) amino acids are interaction-prone and tend to elevate *R*_2_. (The original SeqDYN also has an option of applying a helix boost; we did not apply this boost here.) We transformed SeqDYN into DIRseq by implementing four changes.

First, we reassigned four *q* parameters. We lowered the *q* values of the three long aliphatic amino acids, Leu, Ile, and Met, to the *q* value of a short aliphatic amino acid, Val, because long aliphatic amino acids primarily participate in hydrophobic interactions, which may be less important for stabilizing the binding of a small molecule in sites largely exposed to water. At the same time, we increased the *q* value of Asp to be the same as that of Glu, to increase the role of Asp’s electrostatic interactions. Both the downgrade of hydrophobic interactions and the upgrade of electrostatic interactions were motivated by observations on the drug binding to α-synuclein in MD simulations by Robustelli et al. ^13^. The modified set of *q* parameters is displayed in Figure S1b.

Second, the original *b* value, 0.0316, corresponds to a correlation half-length, *b*^−1/2^, of 5.6 residues. Given the small size of drug molecules, we increased the *b* value to 0.3, corresponding to a correlation half-length of 1.8 residues. That is, we expect a drug molecule to interact with 3 to 4 residues at a time. Third, the SeqDYN-predicted *R*_2_ profile, capturing experimental observations ^36^, falls off at both termini, because no residues beyond the termini are present to provide a contributing factor to *R*_2_. However, whereas intra-IDP interactions experience such a terminal effect, IDP-drug interactions do not. To eliminate the terminal effect, we padded the original IDP sequence by a stretch of 12 Gln residues at each terminus. Gln was selected because its *q* value is at the middle of the 20 *q* values (Figure S1b).

Lastly, we converted high and low *R*_2_ values into high and low drug-interacting propensities, respectively, using a sigmoid function

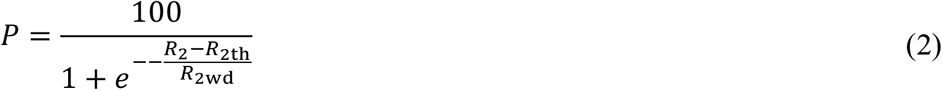

where the midpoint *R*_2th_ and width *R*_2wd_ of the transition region are determined by the mean (*m*) and standard deviation (SD) of the *R*_2_ values over the entire sequence. Specifically,

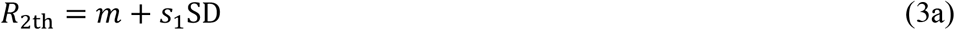

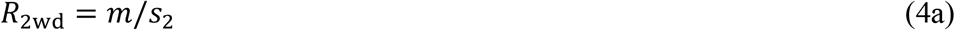

We chose *s*_1_ to be 1.5 and *s*_2_ to be 14.0. On the DIRseq web server, users can either keep these default values for *b, s*_1_, and *s*_2_, or enter values of their choice. There is also a link for users to download the source code. The values of *P* range from 0 to 100; *P* = 50 when *R*_2_ is at the “threshold” value *R*_2th_. Residues with *P* ≥ 50 are predicted to be drug-interacting, but an isolated residue with *P* ≥ 50 is discarded, as it can be reasonably expected that interactions with at least two consecutive (or nearby) residues are needed to generate sufficient drug binding stability.

### Detailed assessment of DIRseq on four IDPs

Drug-interacting residues in four IDPs or intrinsically disordered regions (IDRs) have been thoroughly characterized by NMR CSP. In Figure 1 (left images), we show these IDPs in a single conformation, with DIRseq propensities rendered in a color spectrum (from low in yellow to high in red); predicted drug-interacting residues (i.e., those with *P* ≥ 50) are shown with sidechains in stick. Drug molecules that bind to these IDPs are also displayed in Figure 1 (right images). We display NMR CSPs as blue bars and DIRseq propensities as a red curve in Figure 2. Unless otherwise noted, we use *m* + 1.5 SD as the threshold for identifying drug-interacting residues, for both CSP and DIRseq; this threshold is indicated by a horizontal dashed line in Figure 2. Experimentally identified drug-interacting residues are further indicated by cyan shading. Below we present a detailed comparison between CSP and DIRseq.

**Figure 2.**
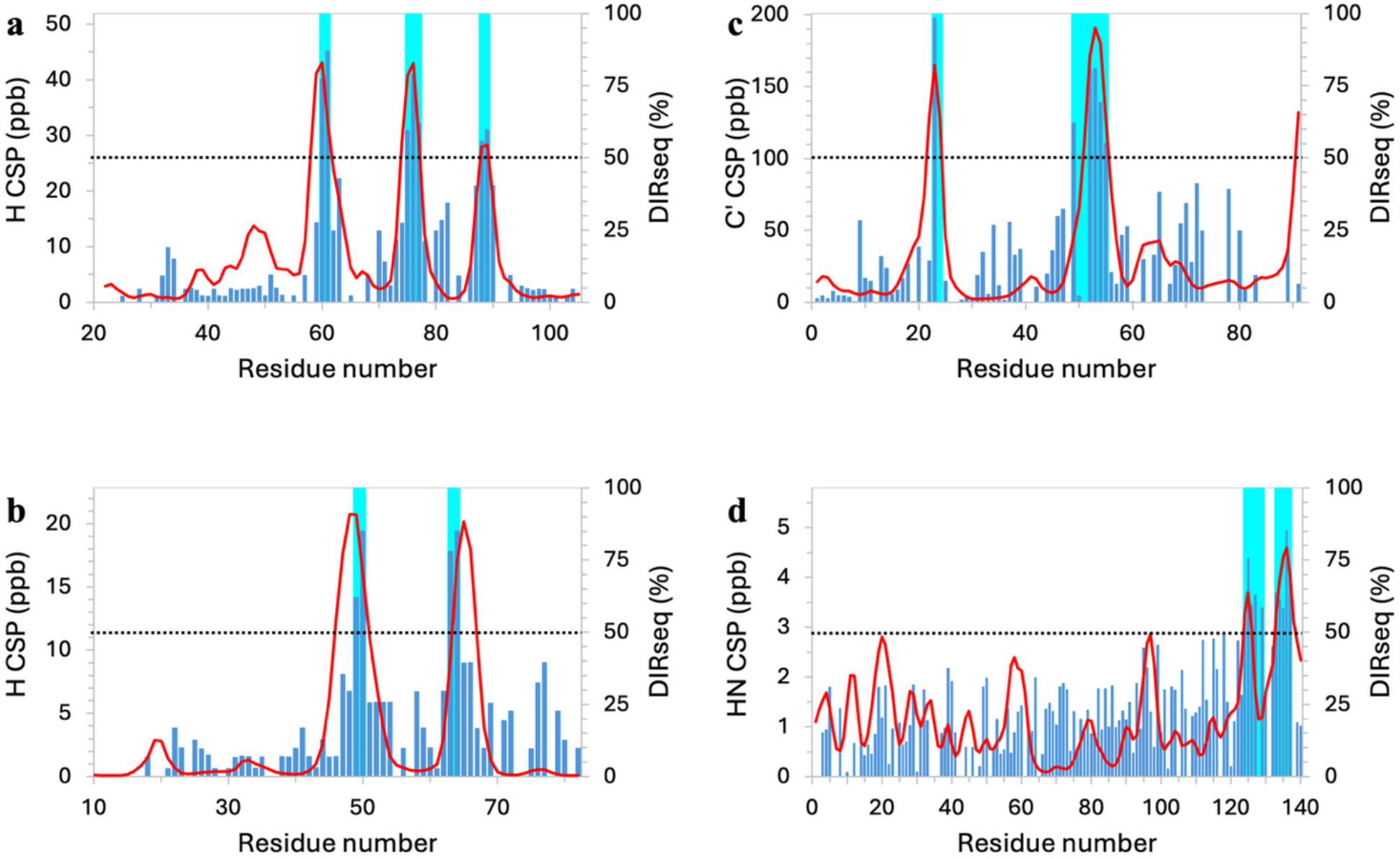
Comparison of DIRseq propensities with NMR CSPs. (a) p27. (b) p21. (c) p53. (d) α-synuclein. CSPs are displayed as blue bars and in units of ppb; CSP-identified drug-interacting residues are indicated by cyan shading. DIRseq predictions are shown as red curves. The ordinate scales are chosen so that the *m* + 1.5 SD threshold for CSP is at the same height as the 50% threshold for DIRseq, indicated by a horizontal dashed line.

The kinase inhibitory domain (residues 22-105) of the cell cycle regulator p27 harbors three aromatic-centered motifs that interact with a group of compounds represented by SJ403 (Figure 1a), as found by Iconaru et al. ^15^. These motifs, W_60_N_61_, E_75_WQ_77_, and Y_88_Y_89_ (Figure 2a, cyan shading), are correctly predicted by DIRseq (Figure 2a, portions of the red curve above the 50% threshold). Specifically, the predicted drug-interacting residues are R_58_KWN_61_, Y_74_EWQ_77_, and Y_88_Y_89_ (Figure 1a, left image). Moreover, the DIRseq propensity profile matches well the CSP profile along the sequence (Figure 2a). The two parameters show a very strong correlation, with a Pearson correlation coefficient (*r*) of 0.82 (Figure S2a).

Iconaru et al. also studied SJ403 binding to a related cell cycle regulator p21. Their CSP data identified the first two corresponding motifs, W_49_N_50_ and F_63_A_64_, but not the third as drug-interacting residues (Figure 2b, cyan shading). DIRseq correctly predicts both the first two motifs above the 50% threshold and the third motif below the threshold (Figure 2b, red curve; Figure 1b, left image). As noted by Iconaru et al., the p21 counterpart to the p27 third motif F_87_YY_89_ is L_76_YL_78_; the two Leu residues in p21 are no match for F_87_ and Y_89_ in p27. For the DIRseq method, Leu has a lower *q* value than Tyr and Phe (Figure S1b). CSP and DIRseq propensity show a strong correlation (*r* = 0.66; Figure S2b).

NMR CSP revealed two motifs, W_23_K_24_ and D_49_IEQWFT_55_, in the N-terminal region of p53 as interacting with EGCG ^12^ (Figure 2c, cyan shading). MD simulations supported these drug-binding sites. DIRseq again correctly predicts these two motifs (Figure 2c, red curve). Specifically, the predicted drug-interacting motifs are L_22_WK_24_ and E_51_QWFT_55_ (Figure 1c, left image). Here also CSP and DIRseq propensity show a strong correlation (*r* = 0.67; Figure S2c).

Using the *m* + 1.5 SD threshold, CSPs of Robustelli et al. ^13^ select residues A_124_YEMPS_129_ and Y_133_QDYE_137_ of α-synuclein as interacting with the drug Fasudil (Figure 2d, cyan shading). These two C-terminal motifs are also identified by DIRseq, with residues A_124_YE_126_ and Y_133_QDYEP_138_ above the 50% threshold (Figure 2d, red curve; Figure 1d, left image). There is a moderate correlation (*r* = 0.51; Figure S2d) between CSP and DIRseq propensity. Robustelli et al. further characterized the IDP-drug interactions by all-atom simulations, using both full-length α-synuclein and a C-terminal fragment (residues 121-140). Further simulations of the C-terminal fragment binding with additional compounds led to a compound known as Ligand 47 with a somewhat higher affinity than Fasudil. The CSP profiles elicited by Fasudil and Ligand 47 are similar, though with larger amplitudes around Y_125_ and Y_136_ by the latter; their correlation coefficient is 0.78 (Figure S3a). Because DIRseq does not consider any information about the drug molecule, this correlation coefficient between CSPs elicited by two different compounds might be viewed as an upper bound of what can be achieved for the correlation between CSP and DIRseq propensity.

### Dependences of DIRseq prediction accuracies on model parameters

We now use the CSP data of the above four IDPs to assess the dependences of DIRseq prediction accuracies on model parameters (Table S2). We use two complementary measures for accuracy: the Pearson correlation between CSP and drug-interacting propensity and the difference between true positive (TP) and false positive (FP). The four IDPs have a total of 31 CSP-identified drug-interacting residues. Recall that DIRseq uses a sigmoid function to convert the *R*_2_ output of SeqDYN into drug-interacting propensities. Using the original 20 *q* parameters along with the present values for *b, s*_1_, and *s*_2_, the predictions are already pretty reasonable: the correlations for the four IDPs range from 0.46 to 0.79 (sum = 2.43) and TP outnumbers FP by 7 residues. Indeed, this initial success was anticipated ^33,34^ and validated the premise of DIRseq, i.e., intra-IDP interactions that elevate *R*_2_ also tend to mediate drug interactions. As stated above, we tweaked four of the 20 *q* parameters to arrive at the final DIRseq; now the *r* sum increases to 2.66 and TP – FP increases to 15.

Examining the four *q* parameter changes one at a time, the downgrade of *q* for Leu increases both *r* sum (to 2.53) and TP – FP (to 16) relative to the counterparts with the original SeqDYN *q* parameters. In comparison, the downgrade of *q* for Ile does not affect *r* sum and yields a small increase in TP – FP (to 10), the upgrade of *q* for Asp yields small increases in both *r* sum (to 2.52) and TP – FP (to 8), but the downgrade of *q* for Met actually decreases both *r* sum (to 2.37) and TP – FP (to 6). The latter result shows that we made these parameter tweaks not solely for increasing accuracy but for physical reasons, i.e., to reduce the role of hydrophobic interactions but elevate the role of electrostatic interactions in IDP-drug binding, as suggested by MD simulations.

We also tested alternative models for *q* parameters. Given the prominence of aromatic amino acids in drug binding as revealed by CSP, we wondered how a model that solely emphasizes aromatic amino acids would perform. To that end, we tested a model with only two different *q* values for the 20 amino acids: a high value (same as that of Trp in SeqDYN) for all three aromatic amino acids (Trp, Tyr, and Phe), and a low value (same as that of Gly in SeqDYN) for all other amino acids. This is similar to a sticker-spacer model for simulating liquid-liquid phase separation ^37^. This aromatic model achieves the same *r* sum, 2.66, as DIRseq, but its FP outnumbers its TP, such that TP – FP = -1. Tesei et al. ^38^ parameterized a coarse-grained force field to simulate liquid-liquid phase separation. Their *λ* “stickiness” parameters have a good correlation with the SeqDYN original *q* parameters ^39^. We replaced the *q* parameters by these *λ* parameters (after scaling to the same numerical range as the *q* parameters); the resulting model has both a low *r* sum (1.21) and a low TP – FP (−21). This outcome suggests that, unlike liquid-liquid phase separation where stickiness is the main drive, drug binding is more selective in the type of intermolecular interactions. Lastly, we tested a model where an average hydropathy scale (compiled by Tesei et al. ^40^) was used in place of the *q* parameters; the resulting model has very little predictive value (*r* sum = 0.44 and TP – FP = -20). This last outcome is in line with our downgrade of hydrophobic interactions in DIRseq. The test results from all these alternative models indicate that the *q* parameters in the final DIRseq model capture the appropriate balance among aromatic, electrostatic, and hydrophobic interactions in IDP-drug binding.

In addition to the 20 *q* parameters, DIRseq has three other parameters. The *b* parameter determines the number of residues that simultaneously interact with a drug molecule. In Table S2, we list the performance measures when *b* is varied. These results show that our final choice, 0.3, for *b* is optimal for both *r* sum and TP – FP. The corresponding number of residues, 3 to 4, that simultaneously interact with a drug molecule fits with the narrow range of drug molecule sizes in the present study (Table S2; molecular weights: 360 ± 130 Da). The two last parameters, *s*_1_ and *s*_2_, are in the sigmoid function that converts the *R*_2_ output into drug-interacting propensities. *s*_1_ sets the *R*_2_ threshold for labeling a residue as drug-interacting. Again, our final choice, corresponding to *m* + 1.5 SD for the threshold, achieves an optimum for *r* sum and TP – FP. A lower threshold leads to a high FP and also a slight deterioration in *r* sum, whereas a higher threshold leads to a low FP and possibly a minuscule improvement in *r* sum. *s*_2_ controls the sharpness of the sigmoid function in the transition region, and affects *r* sum but not TP – FP. *r* sum increases with increasing *s*_2_; we chose an *s*_2_ value where *r* sum is nearly at the saturation level.

### DIRseq as a complement to CSP for assigning drug-interacting residues

After assessing the achievable accuracy of DIRseq, we now consider an application: its combination with CSP to make robust assignments of drug-interacting residues. We note that the above CSP-identified drug-interacting motifs are all anchored on one or more aromatic residues, and this feature likely contributes to the good performance of DIRseq. When a clear “aromatic signal” is not present, CSP or DIRseq alone may not be able to conclusively identify drug-interacting residues. However, a consensus identification by CSP and DIRseq may be reliable. MD simulations have played such a complementary role to CSP in several studies ^12,13,16,30^.

De Mol et al. ^9^ reported the CSPs of an IDR called AF-1* (residues 142-448) in the androgen receptor elicited by EPI-001 and its stereoisomers including EPI-002; small but reproducible CSPs were found in three subregions, R1 (residues 341-371), R2 (residues 391-414), and R3 (residues 426-446). Conversely, AF-1* caused changes in the ^1^H NMR spectrum of EPI-001 but the peptides corresponding to R1-R3 did not, suggesting that, rather than separately interacting with the three subregions, EPI-001 induces a partial folding of the three subregions. MD simulations captured the partial folding of the R2-R3 fragment (residues 391-446) induced by EPI-002, and underscored the importance of aromatic residues in drug binding ^30^. Basu et al. ^16^ reported the CSPs of the transactivation unit 5 (Tau-5*, residues 336-448) in AF-1* by EPI-001 and a more potent variant, 1aa (Figure 3a and Figure S3b). The contact probabilities of the R2-R3 fragment with 1aa in MD simulations reaffirmed the importance of aromatic residues.

**Figure 3.**
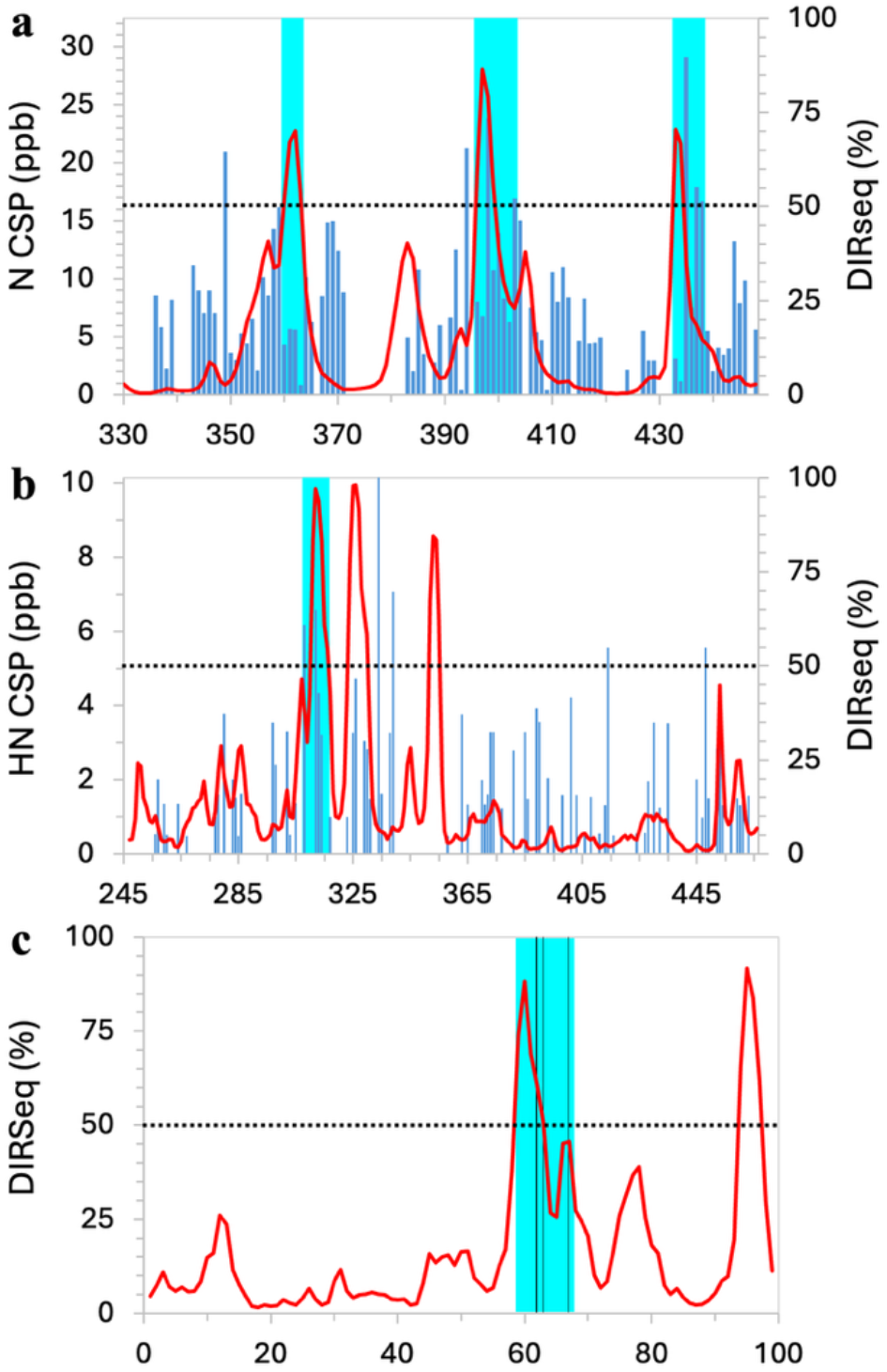
Drug-binding sites identified by combining CSP or mutation data with DIRseq predictions. (a) Tau-5*. (b) NS5A-D2D3. (c) β2 microglobulin. Display items in panels (a, b) have the same meanings as in Figure 2, except that cyan shading indicates consensus identification; in panel (c), vertical lines indicate mutation sites.

De Mol et al. ^9^ and Basu et al. ^16^ were careful not to name any drug-interacting residues based on CSPs. In Figure 3a, we compare the 1aa-elicited CSPs and DIRseq propensities of Tau-5*. In agreement with both NMR studies, DIRseq identifies drug-interacting residues in the middle of each of R1-R3: R_360_DYY_363_, A_396_WAA_399_, and W_433_H_434_. In addition, the latter two motifs showed the highest drug-contact probabilities in separate MD simulations of the R2-R3 fragment _16,30_. For the R2 and R3 subregions, CSPs above the *m* + 1.5 SD threshold are observed at residues downstream of the DIRseq identifications, so we propose to expand the drug-interacting motifs to A_396_WAAAAAQ_403_ and W_433_HTLF_437_. The three putative drug-interacting motifs are indicated as cyan shading in Figure 3a.

Heller et al. ^14^ used ^19^F transverse relaxation measurements to determine the binding affinity of the disordered domains 2 and 3 of the hepatitis C virus NS5A protein (NS5A-D2D3, residues 247-466) for 5-fluorindole. They also measured ^1^H-^15^N CSPs at two ligand concentrations but described them as “nearly undetectable” (Figure 3b). We speculate that the small CSPs may be due to the small size of the ligand, making it difficult to interact with multiple residues simultaneously and thus achieve sufficient binding stability. In any event, CSPs above the *m* + 1.5 SD threshold were largely isolated (and thus appeared to be random), and there was not much overlap between residues having these above-the-threshold CSPs at the two ligand concentrations. The one exception is a motif around W_312_, which had above-the-threshold CSPs at both ligand concentrations, and nearby residues A_308_ and A_313_ also had above-the-threshold CSPs at one of the ligand concentrations. DIRseq predicts the motif P_310_AWARPD_316_ as drug-interacting residues. We propose the expanded motif, A_308_LPAWARPD_316_, as residues interacting with 5-fluorindole (Figure 3b, cyan shading). DIRseq also predicts two more motifs, E_323_SWRRPDY_330_ and R_352_RRR_355_, as drug-interacting residues, which remain to be tested.

The fibrillation of acid-denatured β_2_ microglobulin is inhibited by rifamycin SV ^6^. An aromatic-rich motif, W_60_SFYLLYYTEF_70_, was implicated in the nucleation of fibrillation and also involved in ligand binding, as a triple mutation, F62A/Y63A/Y67A, significantly weakened binding. Low intensities of NMR peaks from residues 58-79 (possibly due to the formation of residual structures) prevented the measurement of CSPs. DIRseq predicts D_59_WSFY_63_ as drug-interacting residues. Combined with the mutational data of Woods et al. ^6^, we propose the expanded motif, D_59_WSFYLLYY_67_, as the major drug-binding site (Figure 3c, cyan shading). DIRseq also predicts an additional motif, K_94_WDR_97_, as drug-interacting residues.

The aggregation of human islet amyloid polypeptide (hIAPP; 37 residues) is inhibited by the small molecule YX-I-1 ^26^. CSPs elicited by this molecule were small (Figure 4a). In addition, CSPs of short IDPs may not exhibit strong disparities because amino acids may be too well mixed along the sequence or drug binding may induce a conformational shift. We thus reduce the threshold for identifying drug-interacting residues to *m* + 1.0 SD when the number of residues is ≤ 50. With this threshold, three residues are identified by CSP as drug-interacting residues: R_11_ and V_17_H_18_. In comparison, DIRseq identifies T_9_QRLA_13_ and F_15_ as drug-interacting residues, which partially overlap with the CSP identifications. Combining the two types of data, we propose T_9_QRLANFLVH_18_ as the primary drug-binding site (Figure 4a, cyan shading). We note that this motif is also prone to α-helix formation ^41^.

**Figure 4.**
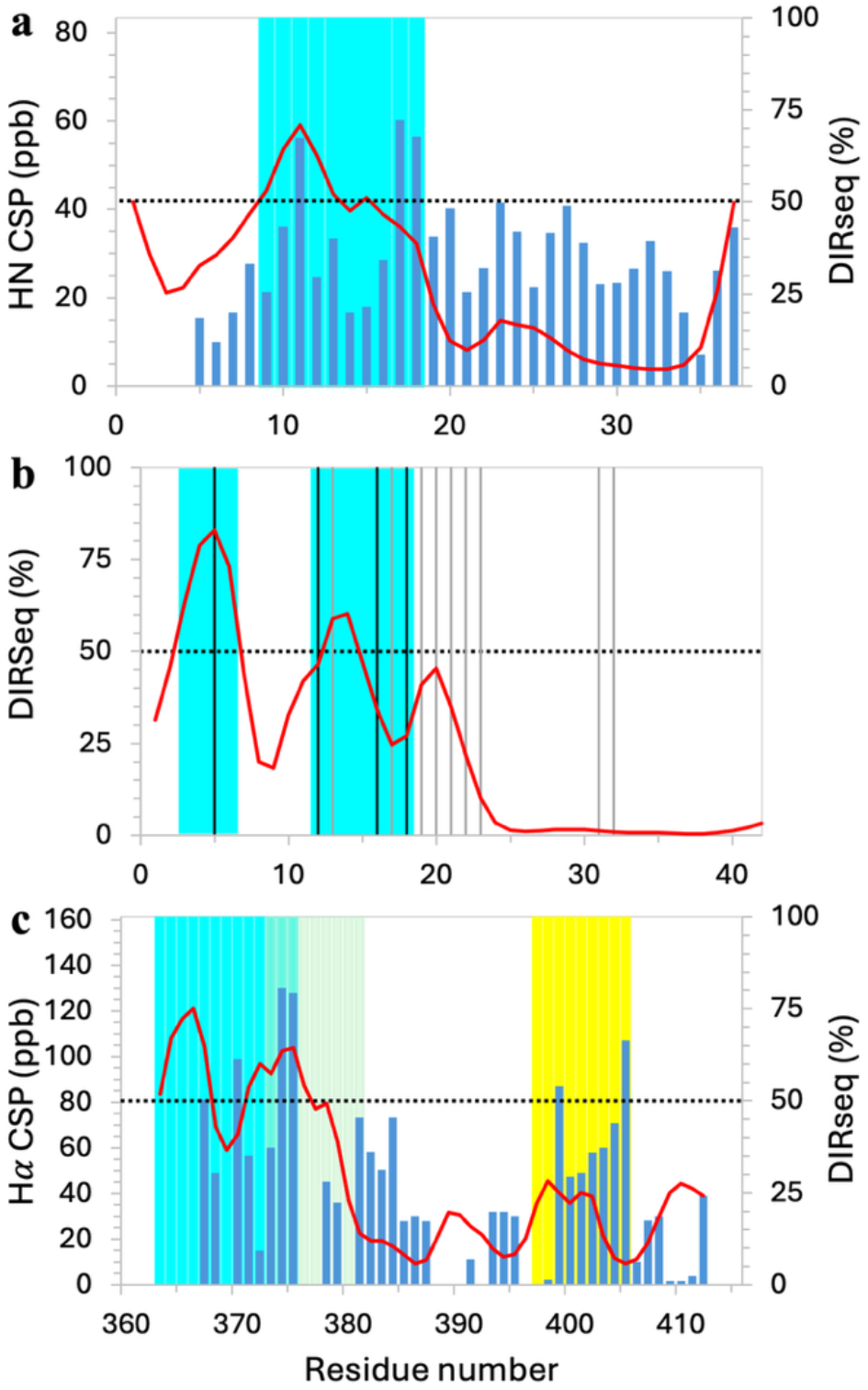
Drug-binding sites identified by combining CSP or mutation data with DIRseq predictions. (a) hIAPP. (b) Aβ42. (c) c-Myc. Display items have the same meanings as in Figure 2, but with the following exceptions. (1) In panel (b), vertical lines indicate residues with prominent CSPs; those accompanied by NMR peak broadening have their vertical lines in dark color. (2) In panel (c), three CSP-identified drug-interacting regions are indicated by cyan, olive, and yellow shading. (3) The threshold for identifying drug-interacting residues is lowered to *m* + 1.0 SD.

Ono et al. ^7^ acquired ^1^H-^15^N heteronuclear single quantum coherence spectra of the 42-residue amyloid-β (Aβ42) in the absence and presence of the oligomerization-blocking compound myricetin. CSPs were not calculated, but chemical shift movements were most pronounced at R_5_, V_12_H_13_, K_16_LVFFAED_23_, and I_31_I_32_ (vertical lines in Figure 4b). In addition, NMR cross peaks suffered broadening upon ligand binding at four of these residues: R_5_, V_12_, K_16_, and V_18_ (dark vertical lines in Figure 4b), implicating elevated probabilities of ligand interactions. DIRseq predicts E_3_FRH_6_ and H_13_H_14_ as drug-interacting residues. Combined with the NMR data, we propose E_3_FRH_6_ and V_12_HHQKLV_18_ as the primary ligand-binding sites (Figure 4b, cyan shading). Aβ42 CSPs elicited by several other compounds were also widely distributed over the sequence, such that Iwaya et al. ^11^ “failed to identify” drug-interacting residues, implicating a conformational shift.

Hammoudeh et al. ^17^ identified three distinct drug binding sites in a C-terminal IDR (residues 363-412) of the oncoprotein c-Myc. This region is disordered on its own but forms a helix-loop-helix structure upon heterodimerization with Max. These authors measured the CSPs of three overlapping fragments (residues 363-381, 370-409, and 402-412) each in the presence of a single compound and of the full IDR in the presence of all three compounds. Using 20 parts per billion (ppb) as the threshold, Hammoudeh et al. named residues R_366_RNELKRSFF_375_, F_375_ALRDQIPELE_385_, and Y_402_ILSVQAE_409_ as the binding sites for 10074-G5, 10074-A4, and 10058-F4, respectively. We present their CSP data for the full IDR in Figure 4c. Using the *m* + 1.0 SD threshold, only six residues are identified as drug-interacting residues: R_367_, L_370_, F_374_F_375_, A_399_, and S_405_. DIRseq predicts two motifs, E_363_RQRR_367_ and K_371_RSFFA_376_, that overlap with the first two sites identified by CSP; a third motif, K_397_KATAY_402_, that corresponds to the third CSP-identified site has moderate drug-interacting propensities. Combining CSP and DIRseq, we revise the three drug-binding sites to be E_363_RQRRNELKRSFF_375_, S_373_FFALRDQI_381_, and K_397_KATAYILS_405_ (Figure 4c, cyan, olive, and yellow shading, respectively). In addition to 10074-G5, 10074-A4, and 10058-F4, many compounds bind to these three sites ^27,42-44^.

## Discussion

We have presented the first sequence-based method, DIRseq, to predict drug-interacting residues of IDPs. Assessment against NMR CSP demonstrates the accuracy of DIRseq. Drug-binding motifs are anchored on one or more aromatic residues, for forming π-π interactions with drug-like molecules that are rich in aromatic groups. The success of DIRseq comes without any specific information on the drug molecules, suggesting that IDPs may have a relatively simple sequence code for drug binding.

The notion that drug-interacting residues may be agnostic to the molecular details of drug compounds is supported by the fact that the same drug can bind to different IDPs. For example, in addition to p53 ^12^ (Figure 2c), the polyphenol EGCG also binds to many other IDPs, including α-synuclein ^45^, hIAPP ^46,47^, Aβ40 ^21^ and Aβ42 ^45^, tau ^48^, and merozoite surface protein 2 ^49^. Likewise, 10074-G5 binds to c-Myc ^17^ (Figure 4c) but also to Aβ42 ^10^. On the other hand, c-Myc represents a case where different compounds bind to distinct sites on a single IDP ^17^. A related example is presented by p27, where SJ403 typifies a group of compounds that share the same three binding sites (Figure 2a). Another group of compounds, typified by SJ710, binds only to the third site. Chemically, the presence of nitrogen atoms in the rings of SJ403 enhances its aromaticity and thus strengthens π-π interactions; in addition, the electronegative groups of SJ403 project into different directions, making it less restricted when forming electrostatic interactions (Figure S4). These features may explain why SJ403 can bind to all three sites whereas SJ710 can bind only to the third site, F_87_YYR_90_, where three consecutive aromatic residues followed by a basic residue ensure that SJ710 can form both π-π and electrostatic interactions. When more data for multiple drugs binding to a single IDP become available, it will be important to use such data to train the next generation of DIRseq where the parameters are drug-specific. As a simple example, the number of residues that can simultaneously bind a drug molecule may grow with the latter’s size; this dependence can be modeled by making the parameter *b* dependent on drug molecule size. The drug molecules studied in the present work have molecular weights of 360 ± 130 Da and thus span a relatively narrow size range.

We have illustrated the combination of DIRseq with NMR CSP to make robust identifications of drug-binding sites in IDPs. Indeed, CSP, MD simulations, and DIRseq are three orthogonal approaches that have great potential in complementing each other, not only for identifying drug-binding sites but also for elucidating the roles of amino acids, their sequence context, and different types of noncovalent interactions in forming such sites. DIRseq offers fast speed and a simple, direct link between sequence motif and propensity for drug binding. As more data from CSP and MD simulations become available, DIRseq can be tuned to make even better predictions. For example, it may be feasible to develop DIRseq versions specific to different pharmacophores.

Another application of DIRseq is to define IDP fragments for in-depth study by MD simulations, as shorter constructs both enable the use of a smaller simulation box and reduce the size of the conformational space. For example, based on CSP data and initial MD simulations of fill-length of α-synuclein, Robustelli et al. ^13^ chose a 20-residue C-terminal fragment for simulations of binding with additional compounds, leading to the identification of Ligand 47 as a stronger binder than the original Fasudil. Similarly, based on CSP data from longer constructs of the androgen receptor, both Zhu et al. ^30^ and Basu et al. ^16^ chose the 56-residue R2-R3 fragment for MD simulations of drug binding. DIRseq can now play a similar role in selecting fragments for MD simulations when CSP data are unavailable. Longer constructs may also present challenges such as resonance assignments to NMR experiments, so well-chosen fragments guided by DIRseq can also benefit NMR studies.

Lastly, virtual screening has been conducted against conformational ensembles of IDPs ^24,50^; drug-binding sites predicted by DIRseq can be used to guide such screening. As a simple illustration, we present poses of EGCG generated by screening against the two DIRseq-predicted binding sites in p53 in Figure 5. As IDPs sample a vast conformational space, knowledge of the binding site can drastically reduce the computational cost. The subset of conformations that generate high docking scores for a given drug at the known site can also provide insight into the mechanism of drug action.

**Figure 5.**
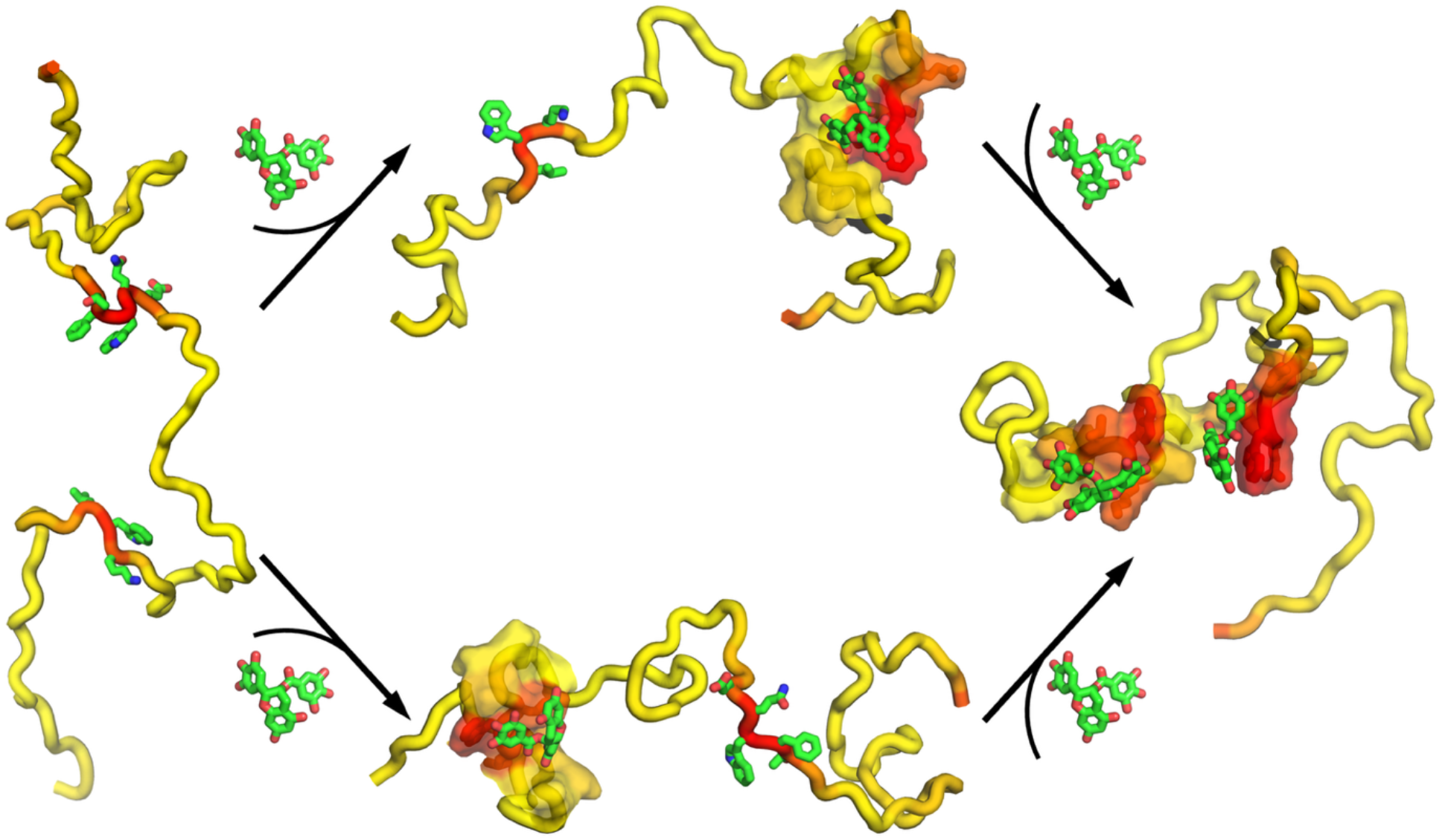
Poses of p53-bound EGCG generated by docking.

### Computational Methods

The sequences of the IDP studied here and the drugs that bind to them are listed in Table S1. All DIRseq predictions were obtained using the web server at https://zhougroup-uic.github.io/DIRseq/. Conformations of IDPs were generated using the TraDES method ^51^.

Docking of EGCG onto p53 was performed via the SwissDock web server at https://www.swissdock.ch/ ^52^ utilizing the Autodock Vina docking engine ^53^. The SMILES string for EGCG from PubChem (https://pubchem.ncbi.nlm.nih.gov/; CID 65064) and several conformations of p53 were used as input. A cubic region (13 to 20 Å in side length) around the center of each drug-interacting residue was selected for docking.

## Acknowledgment

This work was supported by National Institutes of Health Grant GM118091.

## References

(1) Joshi, P.; Vendruscolo, M. In Intrinsically Disordered Proteins Studied by NMR Spectroscopy; Felli, I. C.;Pierattelli, R., Eds.; Springer International Publishing: Cham, 2015, DOI:10.1007/978-3-319-20164-1_13 10.1007/978-3-319-20164-1_13.

(2) Saurabh, S.; Nadendla, K.; Purohit, S. S.; Sivakumar, P. M.; Cetinel, S. Fuzzy Drug Targets: Disordered Proteins in the Drug-Discovery Realm. ACS Omega 2023, 8, 9729–9747.

(3) Wang, H.; Xiong, R.; Lai, L. Rational drug design targeting intrinsically disordered proteins. Wiley Interdiscip Rev Comput Mol Sci 2023, 13, e1685.

(4) Uversky, V. N. How to drug a cloud? Targeting intrinsically disordered proteins. Pharmacol Rev 2024, DOI:10.1124/pharmrev.124.00111310.1124/pharmrev.124.001113.

(5) Peterson, J. R.; Bickford, L. C.; Morgan, D.; Kim, A. S.; Ouerfelli, O.; Kirschner, M. W.; Rosen, M. K. Chemical inhibition of N-WASP by stabilization of a native autoinhibited conformation. Nat Struct Mol Biol 2004, 11, 747–755.

(6) Woods, L. A.; Platt, G. W.; Hellewell, A. L.; Hewitt, E. W.; Homans, S. W.; Ashcroft, A. E.; Radford, S. E. Ligand binding to distinct states diverts aggregation of an amyloid-forming protein. Nat Chem Biol 2011, 7, 730–739.

(7) Ono, K.; Li, L.; Takamura, Y.; Yoshiike, Y.; Zhu, L.; Han, F.; Mao, X.; Ikeda, T.; Takasaki, J.-i.; Nishijo, H. et al. Phenolic Compounds Prevent Amyloid β-Protein Oligomerization and Synaptic Dysfunction by Site-specific Binding. J Biol Chem 2012, 287, 14631–14643.

(8) Jin, F.; Yu, C.; Lai, L.; Liu, Z. Ligand clouds around protein clouds: a scenario of ligand binding with intrinsically disordered proteins. PLoS Comput Biol 2013, 9, e1003249.

(9) De Mol, E.; Fenwick, R. B.; Phang, C. T.; Buzón, V.; Szulc, E.; de la Fuente, A.; Escobedo, A.; García, J.; Bertoncini, C. W.; Estébanez-Perpiñá, E. et al. EPI-001, A Compound Active against Castration-Resistant Prostate Cancer, Targets Transactivation Unit 5 of the Androgen Receptor. ACS Chem Biol 2016, 11, 2499–2505.

(10) Heller, G. T.; Aprile, F. A.; Michaels, T. C. T.; Limbocker, R.; Perni, M.; Ruggeri, F. S.; Mannini, B.; Löhr, T.; Bonomi, M.; Camilloni, C. et al. Small-molecule sequestration of amyloid-β as a drug discovery strategy for Alzheimer’s disease. Sci Adv 2020, 6, eabb5924.

(11) Iwaya, N.; Goda, N.; Matsuzaki, M.; Narita, A.; Shigemitsu, Y.; Tenno, T.; Abe, Y.; Hoshi, M.; Hiroaki, H. Principal component analysis of data from NMR titration experiment of uniformly (15)N labeled amyloid beta (1-42) peptide with osmolytes and phenolic compounds. Arch Biochem Biophys 2020, 690, 108446.

(12) Zhao, J.; Blayney, A.; Liu, X.; Gandy, L.; Jin, W.; Yan, L.; Ha, J. H.; Canning, A. J.; Connelly, M.; Yang, C. et al. EGCG binds intrinsically disordered N-terminal domain of p53 and disrupts p53-MDM2 interaction. Nat Commun 2021, 12, 986.

(13) Robustelli, P.; Ibanez-de-Opakua, A.; Campbell-Bezat, C.; Giordanetto, F.; Becker, S.; Zweckstetter, M.; Pan, A. C.; Shaw, D. E. Molecular Basis of Small-Molecule Binding to alpha-Synuclein. J Am Chem Soc 2022, 144, 2501–2510.

(14) Heller, G. T.; Shukla, V. K.; Figueiredo, A. M.; Hansen, D. F. Picosecond Dynamics of a Small Molecule in Its Bound State with an Intrinsically Disordered Protein. J Am Chem Soc 2024, 146, 2319–2324.

(15) Iconaru, L. I.; Ban, D.; Bharatham, K.; Ramanathan, A.; Zhang, W.; Shelat, A. A.; Zuo, J.; Kriwacki, R. W. Discovery of Small Molecules that Inhibit the Disordered Protein, p27Kip1. Sci Rep 2015, 5, 15686.

(16) Basu, S.; Martínez-Cristóbal, P.; Frigolé-Vivas, M.; Pesarrodona, M.; Lewis, M.; Szulc, E.; Bañuelos, C. A.; Sánchez-Zarzalejo, C.; Bielskutė, S.; Zhu, J. et al. Rational optimization of a transcription factor activation domain inhibitor. Nat Struct Mol Biol 2023, 30, 1958–1969.

(17) Hammoudeh, D. I.; Follis, A. V.; Prochownik, E. V.; Metallo, S. J. Multiple Independent Binding Sites for Small-Molecule Inhibitors on the Oncoprotein c-Myc. J Am Chem Soc 2009, 131, 7390–7401.

(18) Lamberto, G. R.; Binolfi, A.; Orcellet, M. L.; Bertoncini, C. W.; Zweckstetter, M.; Griesinger, C.; Fernández, C. O. Structural and mechanistic basis behind the inhibitory interaction of PcTS on α-synuclein amyloid fibril formation. Proc Natl Acad Sci U S A 2009, 106, 21057–21062.

(19) Krishnan, N.; Koveal, D.; Miller, D. H.; Xue, B.; Akshinthala, S. D.; Kragelj, J.; Jensen, M. R.; Gauss, C.-M.; Page, R.; Blackledge, M. et al. Targeting the disordered C terminus of PTP1B with an allosteric inhibitor. Nat Chem Biol 2014, 10, 558–566.

(20) Tatenhorst, L.; Eckermann, K.; Dambeck, V.; Fonseca-Ornelas, L.; Walle, H.; Lopes da Fonseca, T.; Koch, J. C.; Becker, S.; Tönges, L.; Bähr, M. et al. Fasudil attenuates aggregation of α-synuclein in models of Parkinson’s disease. Acta Neuropathol Commun 2016, 4, 39.

(21) Ahmed, R.; VanSchouwen, B.; Jafari, N.; Ni, X.; Ortega, J.; Melacini, G. Molecular Mechanism for the (−)-Epigallocatechin Gallate-Induced Toxic to Nontoxic Remodeling of Aβ Oligomers. J Am Chem Soc 2017, 139, 13720–13734.

(22) Kocis, P.; Tolar, M.; Yu, J.; Sinko, W.; Ray, S.; Blennow, K.; Fillit, H.; Hey, J. A. Elucidating the Aβ42 Anti-Aggregation Mechanism of Action of Tramiprosate in Alzheimer’s Disease: Integrating Molecular Analytical Methods, Pharmacokinetic and Clinical Data. CNS Drugs 2017, 31, 495–509.

(23) Neira, J. L.; Bintz, J.; Arruebo, M.; Rizzuti, B.; Bonacci, T.; Vega, S.; Lanas, A.; Velázquez-Campoy, A.; Iovanna, J. L.; Abián, O. Identification of a Drug Targeting an Intrinsically Disordered Protein Involved in Pancreatic Adenocarcinoma. Sci Rep 2017, 7, 39732.

(24) Ruan, H.; Yu, C.; Niu, X.; Zhang, W.; Liu, H.; Chen, L.; Xiong, R.; Sun, Q.; Jin, C.; Liu, Y. et al. Computational strategy for intrinsically disordered protein ligand design leads to the discovery of p53 transactivation domain I binding compounds that activate the p53 pathway. Chem Sci 2021, 12, 3004–3016.

(25) Iruela, G.; Fernández, A.; Sagar, A.; Carvajal, F. J.; Bernadó, P.; Pons, M. A FRET-Based Biosensor for the Src N-Terminal Regulatory Element. Biosensors 2022, 12, 96.

(26) Xu, Y.; Maya-Martinez, R.; Guthertz, N.; Heath, G. R.; Manfield, I. W.; Breeze, A. L.; Sobott, F.; Foster, R.; Radford, S. E. Tuning the rate of aggregation of hIAPP into amyloid using small-molecule modulators of assembly. Nat Commun 2022, 13, 1040.

(27) Yu, C.; Niu, X.; Jin, F.; Liu, Z.; Jin, C.; Lai, L. Structure-based Inhibitor Design for the Intrinsically Disordered Protein c-Myc. Sci Rep 2016, 6, 22298.

(28) Herrera-Nieto, P.; Pérez, A.; De Fabritiis, G. Small Molecule Modulation of Intrinsically Disordered Proteins Using Molecular Dynamics Simulations. J Chem Inf Model 2020, 60, 5003–5010.

(29) Wang, G.; Zhu, X.; Song, X.; Zhang, Q.; Qian, Z. Melatonin Inhibits hIAPP Oligomerization by Preventing β-Sheet and Hydrogen Bond Formation of the Amyloidogenic Region Revealed by Replica-Exchange Molecular Dynamics Simulation. Int J Mol Sci 2022, 23.

(30) Zhu, J.; Salvatella, X.; Robustelli, P. Small molecules targeting the disordered transactivation domain of the androgen receptor induce the formation of collapsed helical states. Nat Commun 2022, 13, 6390.

(31) Mehta, T.; Goyal, B. Unveiling molecular mechanism underlying inhibition of human islet amyloid polypeptide fibrillation by benzene carboxylic acid-peptide conjugate. J Mol Liq 2024, 416, 126426.

(32) Lama, D.; Vosselman, T.; Sahin, C.; Liaño-Pons, J.; Cerrato, C. P.; Nilsson, L.; Teilum, K.; Lane, D. P.; Landreh, M.; Arsenian Henriksson, M. A druggable conformational switch in the c-MYC transactivation domain. Nat Commun 2024, 15, 1865.

(33) Qin, S.; Zhou, H. X. Predicting the sequence-dependent backbone dynamics of intrinsically disordered proteins. eLife 2024, 12, RP88958.

(34) Muhammedkutty, F. N. K.; MacAinsh, M.; Zhou, H. X. Atomistic molecular dynamics simulations of intrinsically disordered proteins. Curr Opin Struct Biol 2025, 92, 103029.

(35) Dey, S.; MacAinsh, M.; Zhou, H. X. Sequence-Dependent Backbone Dynamics of Intrinsically Disordered Proteins. J Chem Theory Comput 2022, 18, 6310–6323.

(36) Klein-Seetharaman, J.; Oikawa, M.; Grimshaw, S. B.; Wirmer, J.; Duchardt, E.; Ueda, T.; Imoto, T.; Smith, L. J.; Dobson, C. M.; Schwalbe, H. Long-Range Interactions Within a Nonnative Protein. Science 2002, 295, 1719–1722.

(37) Martin, E. W.; Holehouse, A. S.; Peran, I.; Farag, M.; Incicco, J. J.; Bremer, A.; Grace, C. R.; Soranno, A.; Pappu, R. V.; Mittag, T. Valence and patterning of aromatic residues determine the phase behavior of prion-like domains. Science 2020, 367, 694–699.

(38) Tesei, G.; Lindorff-Larsen, K. Improved Predictions of Phase Behaviour of Intrinsically Disordered Proteins by Tuning the Interaction Range. Open Res Eur 2023, 2, 94.

(39) Zhou, H. X.; Kota, D.; Qin, S.; Prasad, R. Fundamental Aspects of Phase-Separated Biomolecular Condensates. Chem Rev 2024, 124, 8550–8595.

(40) Tesei, G.; Schulze, T. K.; Crehuet, R.; Lindorff-Larsen, K. Accurate Model of Liquid– Liquid Phase Behavior of Intrinsically Disordered Proteins from Optimization of Single-Chain Properties. Proc Natl Acad Sci U S A 2021, 118, e2111696118.

(41) Apostolidou, M.; Jayasinghe, S. A.; Langen, R. Structure of alpha-helical membrane-bound human islet amyloid polypeptide and its implications for membrane-mediated misfolding. J Biol Chem 2008, 283, 17205–17210.

(42) Han, H.; Jain, A. D.; Truica, M. I.; Izquierdo-Ferrer, J.; Anker, J. F.; Lysy, B.; Sagar, V.; Luan, Y.; Chalmers, Z. R.; Unno, K. et al. Small-Molecule MYC Inhibitors Suppress Tumor Growth and Enhance Immunotherapy. Cancer Cell 2019, 36, 483-497.e415.

(43) Shirey, R. J.; Hart, J. R.; Sridharan, B.; Novick, S. J.; Turner, L. D.; Zhou, B.; Nielsen, A. L.; Eubanks, L. M.; Ueno, L.; Hixon, M. S. et al. Synthetic fluorescent MYC probe: Inhibitor binding site elucidation and development of a high-throughput screening assay. Bioorg Med Chem 2021, 42, 116246.

(44) Li, Z.; Huang, Y.; Hung, T. I.; Sun, J.; Aispuro, D.; Chen, B.; Guevara, N.; Ji, F.; Cong, X.; Zhu, L. et al. MYC-Targeting Inhibitors Generated from a Stereodiversified Bicyclic Peptide Library. J Am Chem Soc 2024, 146, 1356–1363.

(45) Ehrnhoefer, D. E.; Bieschke, J.; Boeddrich, A.; Herbst, M.; Masino, L.; Lurz, R.; Engemann, S.; Pastore, A.; Wanker, E. E. EGCG redirects amyloidogenic polypeptides into unstructured, off-pathway oligomers. Nat Struct Mol Biol 2008, 15, 558–566.

(46) Young, L. M.; Cao, P.; Raleigh, D. P.; Ashcroft, A. E.; Radford, S. E. Ion Mobility Spectrometry–Mass Spectrometry Defines the Oligomeric Intermediates in Amylin Amyloid Formation and the Mode of Action of Inhibitors. J Am Chem Soc 2014, 136, 660–670.

(47) Meng, F.; Abedini, A.; Plesner, A.; Verchere, C. B.; Raleigh, D. P. The Flavanol (−)-Epigallocatechin 3-Gallate Inhibits Amyloid Formation by Islet Amyloid Polypeptide, Disaggregates Amyloid Fibrils, and Protects Cultured Cells against IAPP-Induced Toxicity. Biochemistry 2010, 49, 8127–8133.

(48) Wobst, H. J.; Sharma, A.; Diamond, M. I.; Wanker, E. E.; Bieschke, J. The green tea polyphenol (−)-epigallocatechin gallate prevents the aggregation of tau protein into toxic oligomers at substoichiometric ratios. FEBS Lett 2015, 589, 77–83.

(49) Chandrashekaran, I. R.; Adda, C. G.; MacRaild, C. A.; Anders, R. F.; Norton, R. S. Inhibition by Flavonoids of Amyloid-like Fibril Formation by Plasmodium falciparum Merozoite Surface Protein 2. Biochemistry 2010, 49, 5899–5908.

(50) Dhar, A.; Sisk, T. R.; Robustelli, P. Ensemble docking for intrinsically disordered proteins. bioRxiv 2025, DOI:10.1101/2025.01.23.634614 10.1101/2025.01.23.634614, doi: 10.1101/2025.1101.1123.634614.

(51) Feldman, H. J.; Hogue, C. W. Probabilistic sampling of protein conformations: new hope for brute force? Proteins 2002, 46, 8–23.

(52) Bugnon, M.; Röhrig, U. F.; Goullieux, M.; Perez, Marta A. S.; Daina, A.; Michielin, O.; Zoete, V. SwissDock 2024: major enhancements for small-molecule docking with Attracting Cavities and AutoDock Vina. Nucl Acids Res 2024, 52, W324–W332.

(53) Eberhardt, J.; Santos-Martins, D.; Tillack, A. F.; Forli, S. AutoDock Vina 1.2.0: New Docking Methods, Expanded Force Field, and Python Bindings. J Chem Inf Model 2021, 61, 3891–3898.

