## Supplementary Tables and Figures for "DIRseq: a method for predicting drug-interacting residues of intrinsically disordered proteins from sequences"

Table S1 List of IDPs and drugs that bind to them

| IDP name | IDP sequence | Drug name (MW in Da) | Drug structure[a] |
|---|---|---|---|
| p27 (UniProt P46527; residues 22-105) | EHPKPSACRNLFGPVDHEELTRDLEKHCRDMEEASQRKWNFDFQNHKPLEGKYEWQEVEKGSLPEFYYRPPRPPKGACKVPAQE | SJ403 (275.3) | 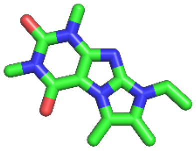 |
| p21 (UniProt P38936; residues 10-82) | QNPCGSKACRRLFGPVDSEQLSRDCDALMAGCIQEARERWNFDFVTETPLEGDFAWERVRGLGLPKLYLPTGP | | |
| p53 (UniProt P04637; residues 1-91) | MEEPQSDPSVEPPLSQETFSDLWKLLPENNVLSPLPSQAMDDLMLSPDDIEQWFTEDPGPDEAPRMPEAAPPVAPAPAAPTPAAPAPAPSW | EGCG (458.4) | 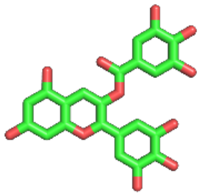 |
| α-synuclein (UniProt P37840; residues 1-140) | MDVFMKGLSKAKEGVVAAAEKTKQGVAEAAGKTKEGVLYVGSKTKEGVVHGVATVAEKTKEQVTNVGGAVVTGVTAVAQKTVEGAGSIAAATGFVKKDQLGKNEEGAPQEGILEDMPVDPDNEAYEMPSEEGYQDYEPEA | Fasudil (291.4) | 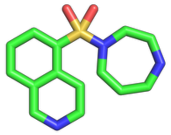 |
| | | Ligand-47 (323.4) | 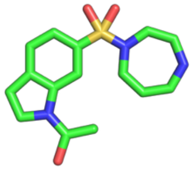 |
| Tau-5* (BMRB 51480; residues 330-448) | AAGSSGTLELPSTLSLYKSGALDEAAAYQSRDYYNFPLALAGPPPPPPPPHPHARIKLENPLDYGSAWAAAAAQCRYGDLASLHGAGAAGPGSGSPSAAASSSWHTLFTAEEGQLYGPC | 1aa (376.8) | 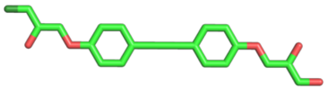 |
| | | EPI-001 (394.9) | 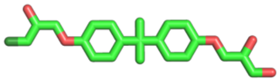 |
| NS5A-D2D3 (residues 247-466, corresponding to UniProt Q99IB8 | SNTYDVDMVDANLLMEGGVAQTEPESRVPVLDFLEPMAEEESDLEPSIPSECMLPRSGFPRALPAWARPDYNPPLVESWRRPDYQPPTVA | 5-Fluoroindole (135.1) | 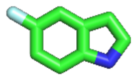 |

| | | | |
|---|---|---|---|
| residues 2223-2442) | GCALPPPKKAPTPPP RRRRTVGLSESTISEA LQQLAIKTFGQPPSS GDAGSSTGAGAAES GGPTSPGEPAPSETGS ASSMPPLEGEPGDPD LESDQVELQPPPQGG GVAPGSGSGSWSTCS EEDDTTVCC | | |
| β2 microglobulin (UniParc UPI0000110347; residues 1-99) | IQRTPKIQVYSRHPAE NGKSNFLNCYVSGF HPSDIEVDLLKNGERI EKVEHSDLSFSKDWS FYLLYYTEFTPTEKD EYACRVNHVTLSQPK IVKWDRDM | Rifamycin SV (697.8) | 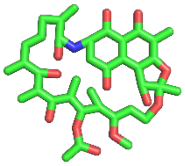 |
| hIAPP (UniParc UPI000002B886; residues 1-37) | KCNTATCATQRLANF LVHSSNNFGAILSSTN VGSNTY | YX-I-1 (435.5) | 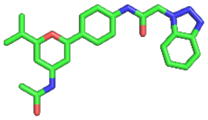 |
| Aβ42 (UniParc UPI00000315E8; residues 1-42) | DAEFRHDSGYEVHH QKLVFFAEDVGSNKG AIIGLMVGGVVIA | Myricetin (320.3) | 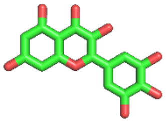 |
| c-Myc (UniProt P01106-1; residues 363-412) | ERQRRNELKRSFFAL RDQIPELENNEKAPK VVILKKATAYILSVQ AEEQK | 10074-G5 (332.3) | 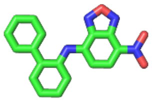 |
| | | 10074-A4 (409.3) | 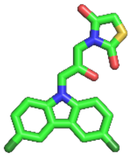 |
| | | 10058-F4 (249.4) | 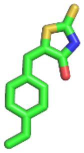 |

[a]Atom color scheme: carbon, green; nitrogen, blue; oxygen, red; sulfur, yellow, fluorine, sky blue; and chlorine, dark green.

Table S2 Dependences of prediction accuracies on model parameters

| | *r* | | | | *r* sum | FP | FN | TP | TP-FP |
|---|---|---|---|---|---|---|---|---|---|
| | p27 | p21 | p53 | α-syn | | | | | |
| *q* parameters | | | | | | | | | |
| seqDYN orig | 0.79 | 0.57 | 0.61 | 0.46 | 2.43 | 12 | 12 | 19 | 7 |
| DIRseq L only | 0.82 | 0.55 | 0.67 | 0.49 | 2.53 | 8 | 7 | 24 | 16 |
| DIRseq I only | 0.79 | 0.62 | 0.56 | 0.46 | 2.43 | 10 | 11 | 20 | 10 |
| DIRseq M only | 0.80 | 0.58 | 0.60 | 0.39 | 2.37 | 11 | 14 | 17 | 6 |
| DIRseq D only | 0.73 | 0.63 | 0.62 | 0.54 | 2.52 | 12 | 11 | 20 | 8 |
| DIRseq | 0.82 | 0.66 | 0.67 | 0.51 | 2.66 | 10 | 6 | 25 | 15 |
| Aromatic | 0.74 | 0.74 | 0.56 | 0.62 | 2.66 | 22 | 10 | 21 | -1 |
| CALDAVOS2 | 0.52 | 0.23 | 0.33 | 0.13 | 1.21 | 27 | 25 | 6 | -21 |
| Avg HPS scale | 0.13 | 0.01 | 0.16 | 0.14 | 0.44 | 23 | 28 | 3 | -20 |
| *b* value | | | | | | | | | |
| 0.0316 | 0.32 | 0.58 | 0.57 | 0.63 | 2.10 | 15 | 11 | 20 | 5 |
| 0.1 | 0.77 | 0.66 | 0.64 | 0.59 | 2.66 | 14 | 11 | 20 | 6 |
| 0.3 | 0.82 | 0.66 | 0.67 | 0.51 | 2.66 | 10 | 6 | 25 | 15 |
| 0.5 | 0.81 | 0.65 | 0.66 | 0.48 | 2.60 | 9 | 7 | 24 | 15 |
| 1 | 0.77 | 0.61 | 0.65 | 0.42 | 2.45 | 7 | 12 | 19 | 12 |
| 3 | 0.68 | 0.51 | 0.58 | 0.32 | 2.09 | 4 | 21 | 10 | 6 |
| $s_1$ value | | | | | | | | | |
| 0.5 | 0.71 | 0.60 | 0.55 | 0.42 | 2.28 | 71 | 2 | 29 | -42 |
| 1 | 0.79 | 0.65 | 0.61 | 0.47 | 2.52 | 35 | 4 | 27 | -8 |
| 1.5 | 0.82 | 0.66 | 0.67 | 0.51 | 2.66 | 10 | 6 | 25 | 15 |
| 2 | 0.82 | 0.63 | 0.71 | 0.55 | 2.71 | 5 | 19 | 12 | 7 |
| 2.5 | 0.80 | 0.58 | 0.72 | 0.58 | 2.68 | 1 | 25 | 6 | 5 |
| $s_2$ value | | | | | | | | | |
| 5 | 0.74 | 0.62 | 0.63 | 0.43 | 2.42 | 10 | 6 | 25 | 15 |
| 10 | 0.79 | 0.65 | 0.65 | 0.48 | 2.57 | 10 | 6 | 25 | 15 |
| 14 | 0.82 | 0.66 | 0.67 | 0.51 | 2.66 | 10 | 6 | 25 | 15 |
| 20 | 0.84 | 0.66 | 0.67 | 0.56 | 2.73 | 10 | 6 | 25 | 15 |

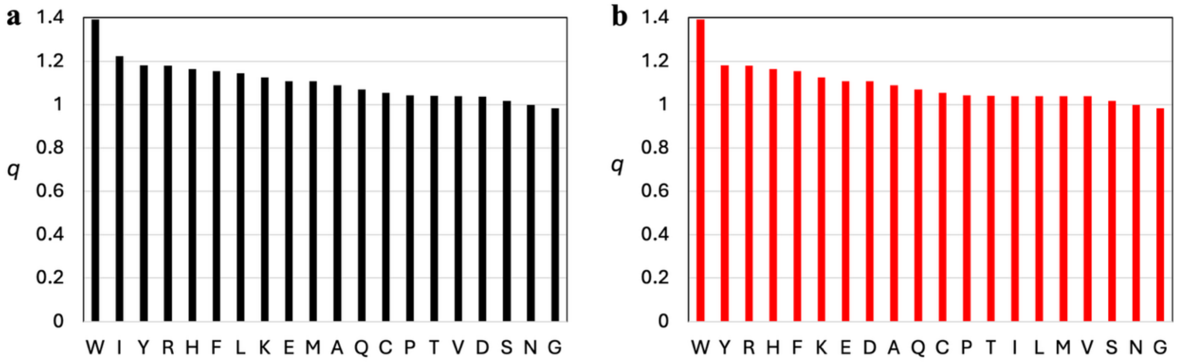


Figure S1. $q$ parameters. (a) Values in the original SeqDYN method (ref. 29). (b) Modified values for DIRseq.

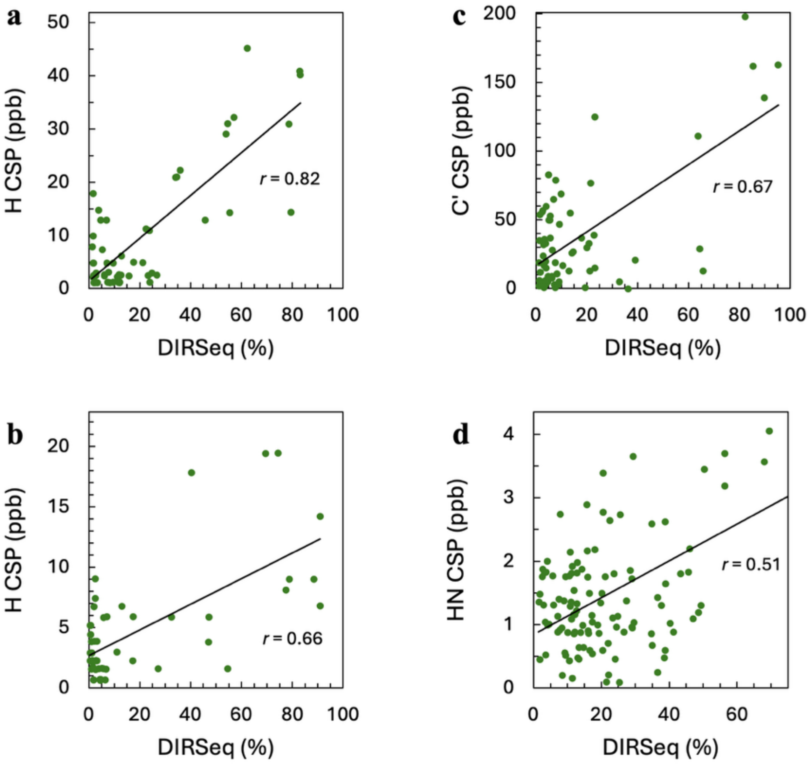


Figure S2. Correlation between NMR CSP and DIRseq propensity. (a) p27. CSP data from ref. 10. (b) p21. CSP data from ref. 10. (c) p53. CSP data from 16. (d) $\alpha$-synuclein. CSP Data from ref. 18.

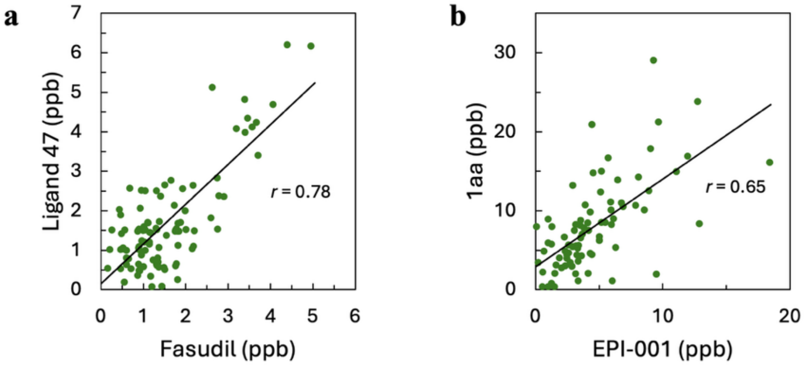


Figure S3. Correlation of CSPs elicited by two different compounds. (a) α-synuclein. Data from ref. 18. (b) Tau-5*. Data from ref. 20.

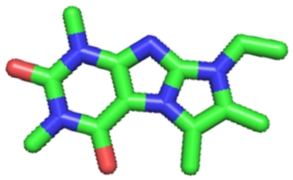

**SJ403**

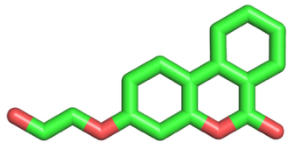

**SJ710**

Figure S4. Two compounds that bind to p27 with different characteristics.
